# deepthought: the microscopy acquisition stack as an object of study

**DOI:** 10.1101/2025.02.25.639997

**Authors:** P.S. Kesavan, Subindev Devadasan, Darshika Bohra

## Abstract

Analysis-in-the-loop microscopy has been demonstrated many times, but it is rarely used outside the laboratories that build it. Each demonstration constructs its own acquisition infrastructure, so little transfers between them, and it has remained unclear which parts of the problem are already solved. In this work, the microscopy acquisition stack was itself treated as the object of study, and was investigated by construction. A minimal stack was built end to end, and four applications were then driven through it as test conditions. These were fixed-cell high-throughput immunofluorescence, live time-lapse imaging of an unsynchronized population, fluorescence anisotropy imaging, and autonomous focus and exposure. Each element of the stack was then classified by how it varied across these applications. Device access, sequencing, data storage and viewing held constant, and mature implementations of each were adopted unchanged. Interpretation, or how an image becomes a set of entities, differed with the application and belongs behind an interface. Two elements had nothing available to adopt and were therefore built. These are a geometric representation of the sample that a plan can traverse, and a representation of a run that yields detected objects rather than images. With those two in place, feedback from analysis into acquisition was ordinary control flow. The system was applied to the DNA damage response, where 22,000 cells were acquired and analyzed without operator intervention, and cells were followed through mitosis over 24 hours in an unsynchronized population without chemical synchronization. One element, targeting, or the choice of where to observe next, varies between applications and remains unabstracted in this implementation. It is identified here as the next requirement.

**Code:** https://github.com/ndsystems/deepthought (acquisition client), https://github.com/ndsystems/hard-link (instrument access server)

## 1 Introduction

The case for coupling image analysis to image acquisition has been made for six decades [Wald, 1966, Herron et al., 1972, Brenner et al., 1976]. When analysis runs during acquisition, the microscope can respond to what it finds, such as skipping empty fields, revisiting interesting ones, correcting focus and exposure, etc. Working demonstrations of this exist [Scherf and Huisken, 2015, Mahecic et al., 2022, André et al., 2023], and recent frameworks have made the underlying plumbing considerably more approachable [Roos et al., 2024, Marin et al., 2024].

Adoption of such systems nevertheless remains limited. The obstacle appears to be structural and not conceptual. Each demonstration is built as a complete stack of its own, assembled for one application, on one instrument, with the feedback path threaded through it by hand. Since each stack is built for a single demonstration, its components are rarely reused by other groups, and the question of which parts of the problem were already solved, and which had to be invented, is seldom addressed directly.

In this work, the stack itself was treated as the object of study. Instead of designing an architecture and then arguing for it, or reviewing the architectures of existing systems, a stack was built in order to study one. The thinnest stack that could run a real experiment end to end was constructed first, and several unrelated applications were then driven through it as test conditions. Each application exercises the same elements of the stack differently, and comparing them shows which elements are properties of microscopy and which are properties of a particular experiment.

This paper reports what that construction showed. The result is a classification and not a framework. Most of the stack is already solved by mature community software and can simply be adopted. Two elements differ legitimately between applications and are better treated as interfaces. For two elements, no implementation was found that could be adopted, and these two were developed as part of this work.

The study also identified an element that was not abstracted here. Targeting, or the decision of where to observe next, varies between applications. One strategy for answering it was adopted, but the element itself was never exposed as an interface, so a different strategy cannot be substituted for it. It is reported below as an open requirement and not as unfinished work.

## 2 Studying the stack by construction

### 2.1 Definitions and classification criterion

A stack can be taken in depth, as the ordered set of elements that must each be answered once for a microscope to produce data. It can also be taken in breadth, as the spread of applications driven through it. A single stack, built minimally, forces every element to be instantiated at once and thereby reveals what the elements are. Driving further applications through it then tests each element against conditions it was not built for. An element that holds constant as applications change can be considered a property of microscopy, while an element that must change with the application is a property of that experiment.

For this comparison to be meaningful, the elements have to be commensurable. A single criterion was used, in which an element of the stack is a question that the stack must answer exactly once per run. If a question can be posed twice at different levels of the system, it is two elements. If it arises in only one application, it is not an element of the stack but an implementation detail within one. This criterion excludes image registration, for instance, which is required by anisotropy imaging and by nothing else. It also excludes the choice of transport protocol, which is not a question the stack answers but a decision about where the stack is divided between processes.

### 2.2 Applications used as test conditions

Four applications were driven through the stack. Fixed-cell high-throughput immunofluorescence requires systematic traversal of a large sample together with per-object intensity measurement across channels. Live time-lapse imaging of an unsynchronized population requires positions to be revisited over many hours with focus maintained between visits. Fluorescence anisotropy imaging requires a single field to be split into two polarization components, registered, and then combined arithmetically. Autonomous focus and exposure require a measurement computed from an image to drive an actuator during the run.

These applications differ in what they demand of the stack, such as coverage in space, sampling in time, optical manipulation within a frame, and feedback onto an actuator. Two of them, being high-throughput immunofluorescence and live time-lapse, were carried to completion and produced the biological results reported below. Anisotropy imaging and autonomous exposure were built only far enough to exercise the elements of the stack and were not carried to a biological result. A fifth application, in which fields are placed by clustering previously detected objects instead of on a grid, was specified and partially implemented, and it is discussed in Section 6.2 because of what it revealed.

### 2.3 Classification of stack elements

Table 1 gives the result. Eight elements were identified, and each falls into one of four categories, the category following from the element’s behaviour across the applications.

**Table 1.**
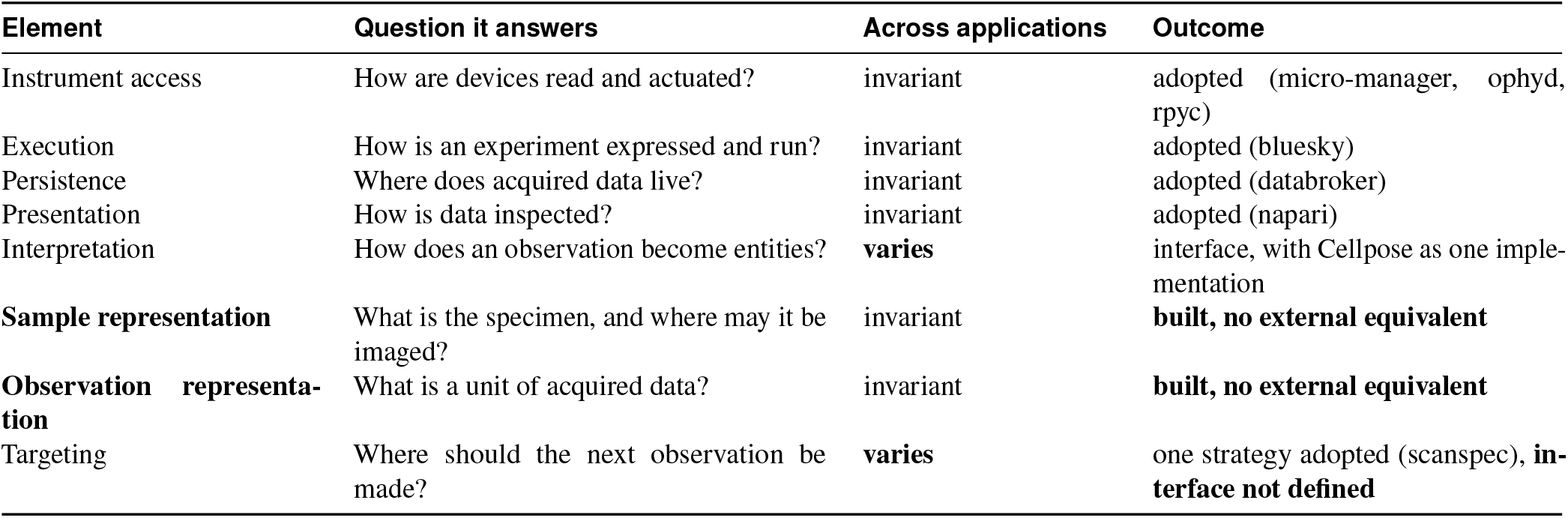
Each element of the stack, classified by its behaviour across the applications tested. Invariant elements with mature external implementations were adopted and not written. Elements that vary between applications can be treated as interfaces. For two invariant elements no implementation was found that could be adopted, and these were built. They are the contribution of this work. One further element, targeting, was identified late and remains unabstracted.

Four elements were invariant across applications and already had mature external implementations, and these were adopted and not written. The gain here is not only in effort saved. The ophyd library supplies an explicit definition of a device, being anything that can be set and read, and bluesky [Allan et al., 2019] supplies an explicit definition of an experiment, being a plan submitted to an engine and not a script that runs. In each case the adopted implementation made explicit a model that the corresponding hand-written code had left implicit.

Two elements vary between applications and are better treated as interfaces. Interpretation is the clear case, since segmenting nuclei from a DAPI channel and dividing a frame into two polarization components share no implementation, although both answer the same question and both must produce a labelled image that downstream code can consume. An interface with interchangeable implementations allows a single stack to serve both. The second varying element, targeting, was not implemented as an interface in this system, for reasons discussed in Section 6.2.

The remaining two elements were invariant across applications with no external equivalent available. These had to be built, and they are where the contribution of any such system can be expected to lie.

### 2.4 Relation between elements and architectural layers

The eight elements of Table 1 group into the four layers of Figure 1. Instrument access forms the device layer, execution forms the plan layer, sample representation and targeting together form the sample layer, and interpretation together with observation representation forms the analysis layer. Persistence and presentation sit outside this grouping, since they consume the data stream and do not produce it.

**Figure 1.**
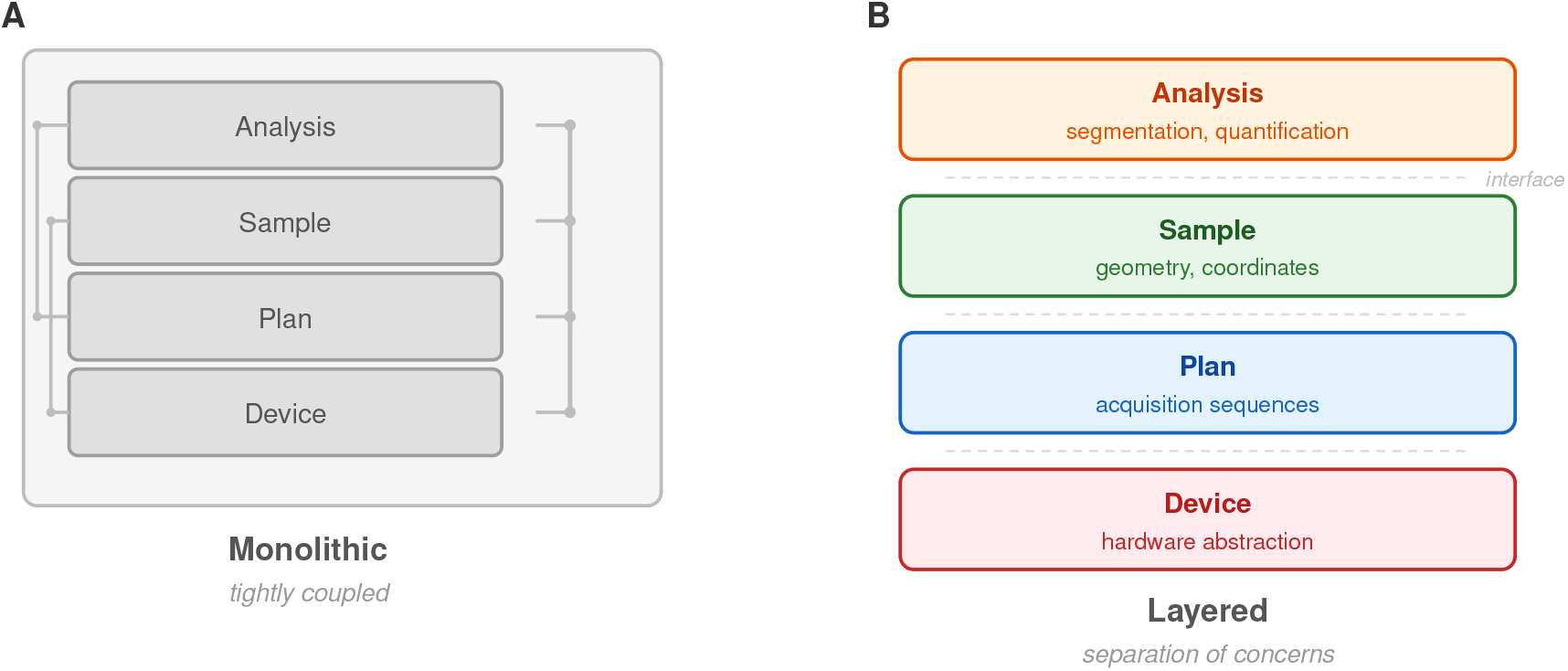
The outcome of the study. Monolithic microscopy software (left) couples device control, experimental logic and data handling into one unit, so that each new application requires the whole stack to be rebuilt. The structure on the right is what the study produced, being the four layers that held constant across every application tested, separated at the boundaries this exposed. These layers are a coarser view of the eight elements given in Table 1, two of which had to be built.

The grouping is coarser than the elements, and one consequence of this matters. In a four-layer view, sample representation and targeting fall into a single layer, since both can naturally be described as belonging to the sample. They behave differently across applications, one being invariant and the other not, so a four-layer view cannot distinguish them. Resolving the stack at the level of elements separates the two, and it was only at that resolution that the omission described in Section 6.2 became visible.

## 3 Components developed in this work

Two elements were invariant across the applications tested and had no implementation that could be adopted. They can be stated as two questions that any acquisition stack has to answer. The first is what a sample is, meaning what the specimen consists of geometrically and where on it imaging is permitted.

The second is what a run produces, meaning whether the output of an acquisition is a set of images or a set of detected objects. Both were built here, and they are described in turn below.

### 3.1 Sample representation, or what a sample is

Every application must answer where on the specimen it is permitted to observe, and the answer can be considered a property of the specimen and not of the instrument or of the experiment. A confocal dish is a circle of known centre and diameter, a coverslip is a rectangle, and a multi-well plate is an array of isolated circles with a known pitch (Fig. 2). Encoding this geometry in an object, instead of in the plan that traverses it, allows a scan to be expressed as the intersection of a traversal pattern with a permitted region. In this system, that intersection keeps acquisition on the glass and away from the plastic periphery of the dish, where refraction degrades image quality, without the plan containing any statement about dishes.

**Figure 2.**
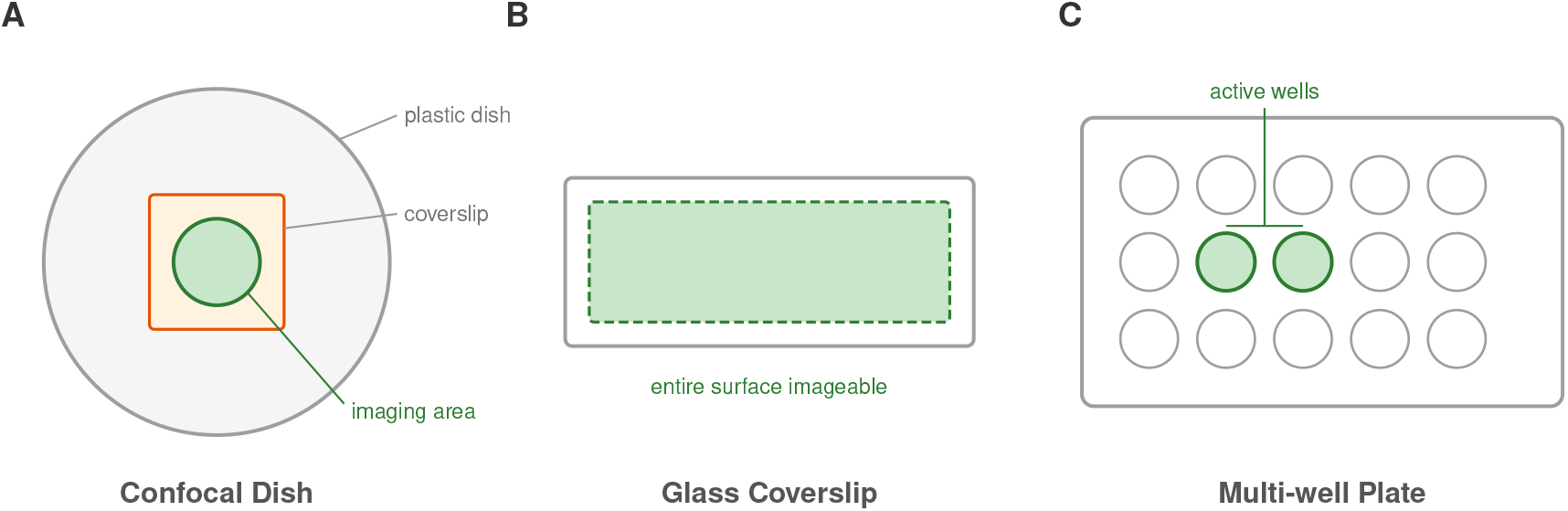
The sample as a geometric object. Common formats, such as confocal dishes, coverslips, multi-well plates, etc., impose distinct constraints on where imaging is possible. Encoding these constraints as properties of a sample object allows plans to traverse any of them without format-specific logic, and allows measurements to be associated with positions on the specimen and not with stage coordinates alone.

The same abstraction serves a second purpose that only became apparent with the live-imaging application. If measurements are associated with positions on the sample and not with raw stage coordinates, positions can be revisited across hours and across sessions, making queries of the form “return to the objects whose signal exceeded a threshold” expressible. Existing projects abstract devices well. We are not aware of one that treats specimen geometry as an object which a plan can traverse, although plate layout is commonly represented as acquisition metadata.

### 3.2 Observation representation, or what a run produces

The second element developed here concerns what a run produces. Under the conventional model, a run yields images, and segmentation is performed afterwards in a separate program. Under the model arrived at here, a run yields detected objects, in that each acquired frame is interpreted as it arrives, and the detected entities accumulate during acquisition together with their positions on the sample, their intensities in each channel, their morphological properties, etc.

Interpreting frames during the run changes what the plan is able to branch on, and this is the reason the closed loop reported here required no dedicated machinery. If the system holds images and the plan needs a decision, something has to bridge the two, and that bridge is the purpose-built code which earlier demonstrations have had to write [Mahecic et al., 2022]. If acquisition already emits entities with positions attached, the plan can branch on the same structure the run is producing, and no bridge is needed (Fig. 3).

**Figure 3.**
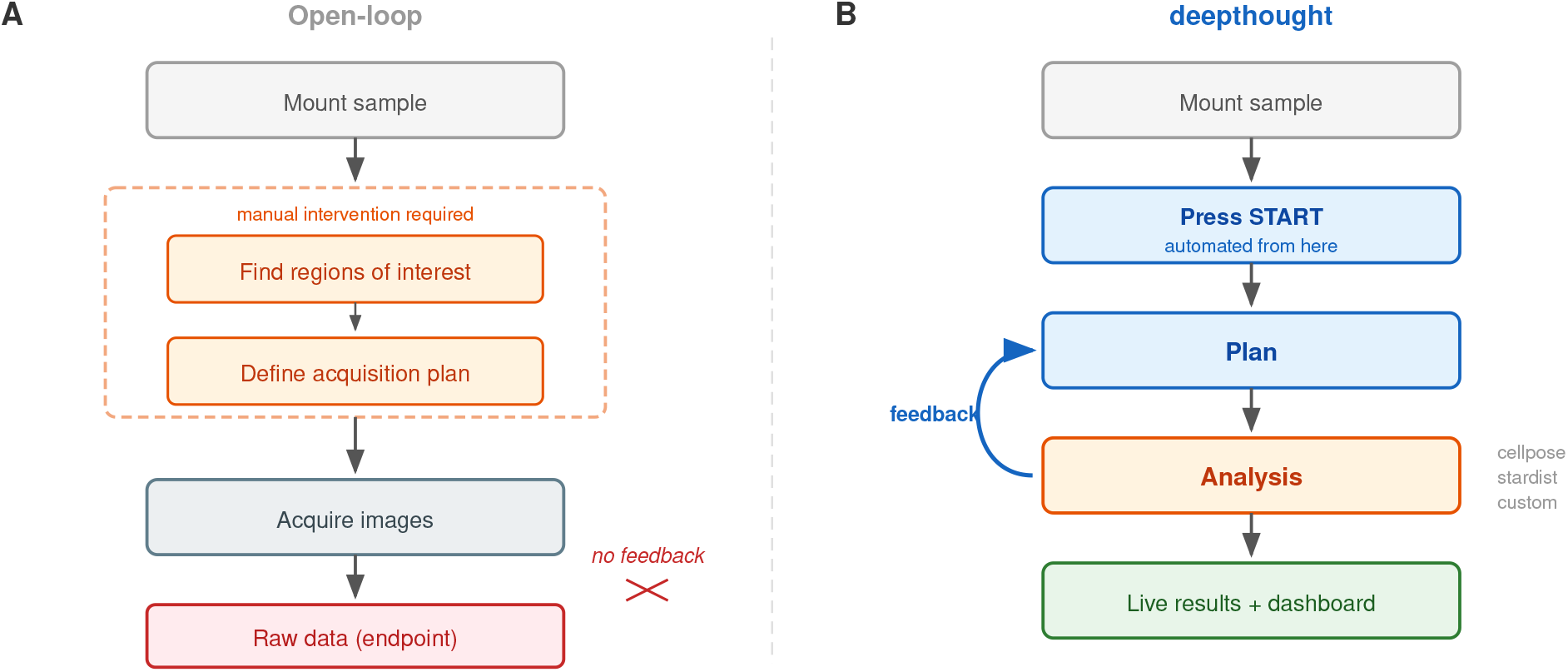
Closed-loop operation as a consequence of representation. In the open-loop arrangement (left), fields are chosen manually, raw data is the endpoint, and analysis occurs after the experiment. In the arrangement reported here (right), interpretation occurs within the run and the plan consumes detected objects. No component mediates between analysis and acquisition, since both operate on the same representation.

## 4 Components adopted from existing software

The remainder of the stack is community software, and it is reported here as a result of the study and not as an implementation detail. Instrument access uses micro-manager [Edelstein et al., 2014] through MMCore, its unified device interface, reached from Python via pymmcore, with devices presented to the rest of the system through ophyd. Execution uses bluesky [Allan et al., 2019], whose RunEngine handles sequencing, interruption, metadata and the emission of documents. Persistence uses databroker, which catalogues those documents and makes a completed or in-progress run addressable. Presentation uses napari [napari contributors, 2019]. Interpretation uses Cellpose [Stringer et al., 2021] as its default implementation. Targeting, where it is answered by a grid, uses scanspec, which represents a traversal as a composable specification and not as nested loops.

Each of these was written by hand in an earlier version of this system before being replaced, so their adoption is itself an outcome of the study.

One decision cuts across these elements without belonging to any of them. Instrument access runs in a separate process from everything else and is exposed over the network (Fig. 4). This arrangement was adopted because instruments are slow to reinitialize, so the process that owns the hardware has to outlive the code being written against it. It was retained because client code can then run on any operating system regardless of what the instrument requires, because a failure in experimental logic cannot disturb the hardware session, and because a single client can hold connections to several instruments at once.

**Figure 4.**
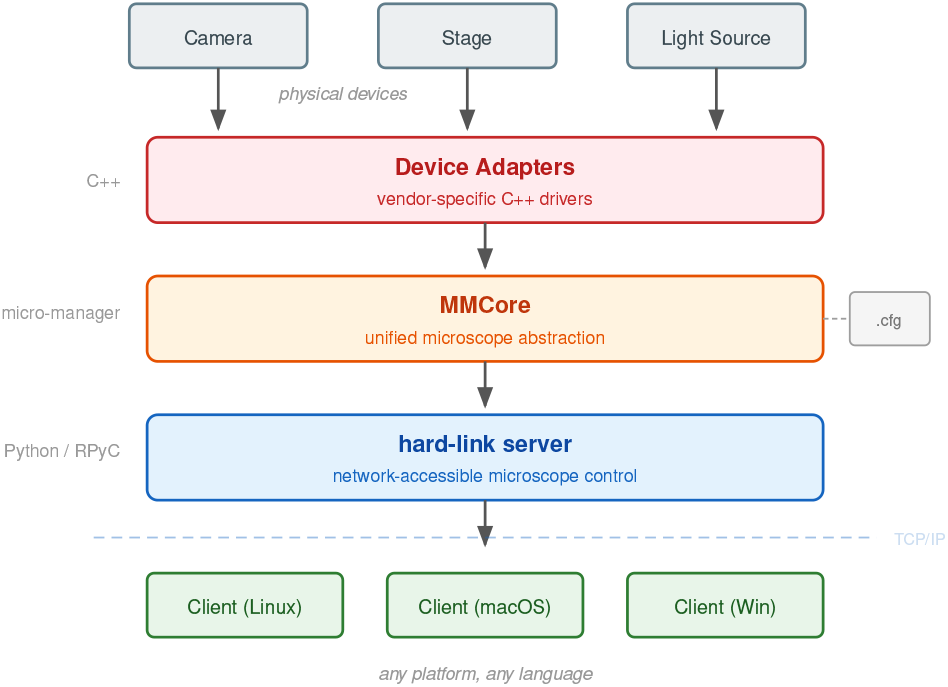
Instrument access across a process boundary. Devices connect through micro-manager adapters to its core interface, which is exposed over TCP/IP by a server process. Since the boundary is a network interface and not a library call, clients are independent of the instrument’s operating system and are free to fail without affecting the hardware session.

The transport crossing this boundary was rewritten once, for a reason that is likely to recur in similar systems. MMCore is a C++ library exposed to Python through generated bindings, and its objects hold live hardware state which cannot be meaningfully copied across a process boundary. A transport built on serialization therefore cannot carry the objects the system exists to manipulate. What is required instead is a transport that proxies remote objects by reference, and rpyc was adopted for this purpose. The sequence here, in which a transport was written by hand, a constraint was discovered, and an existing implementation satisfying that constraint was then adopted, is the same sequence that produced the classification as a whole.

## 5 Results

The system was applied to the DNA damage response in cultured cells. Two experiments are reported, covering unattended acquisition at scale and a rare transition captured in an unsynchronized live population.

### 5.1 Acquisition at scale without an operator

HeLa cells were treated with neocarzinostatin (NCS), a radiomimetic agent inducing double-strand breaks, and stained for *γ*H2AX and phosphorylated Chk1. The plan traversed the sample, and interpretation segmented nuclei from the DAPI channel as frames arrived and extracted per-object intensities in the remaining channels. The system acquired and analyzed 22,000 cells, being 11,000 per condition, with no operator intervention after the run was started.

NCS produced an increase in *γ*H2AX with no detectable change in phosphorylated Chk1 (Fig. 5). The direction of this effect is expected. What is relevant here is that the measurement was made on an unmodified motorized wide-field microscope, at a sample size that is far higher than what is regularly obtained in microscopic investigations of the damage response on such instruments.

**Figure 5.**
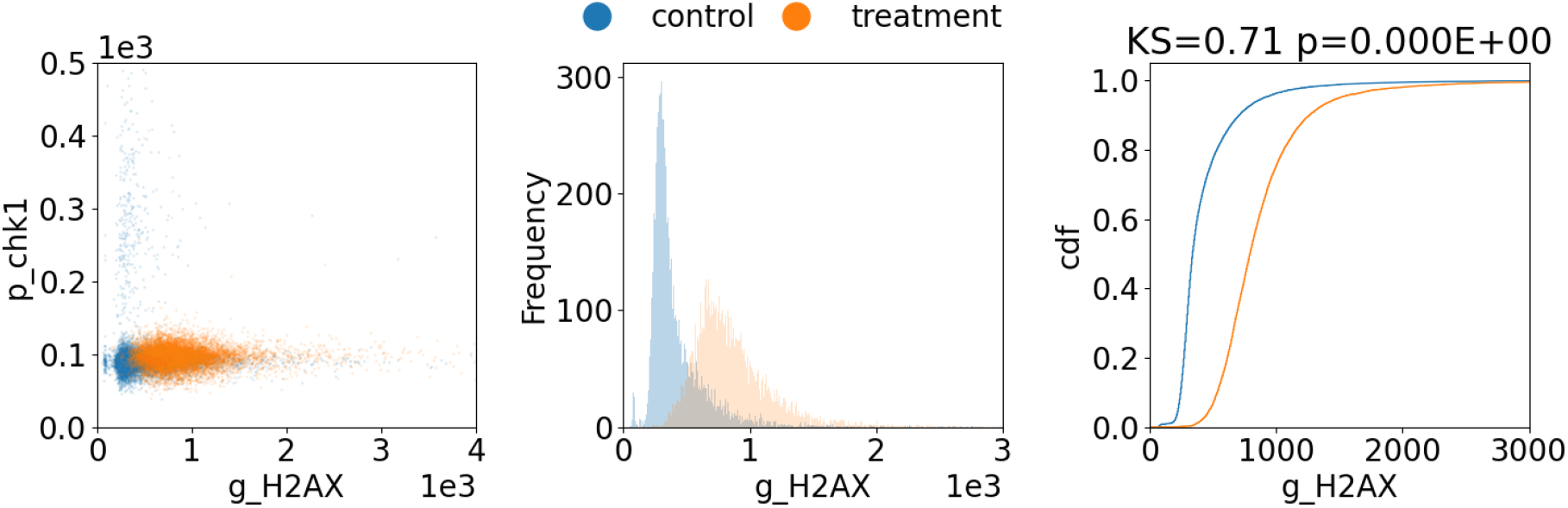
Automated immunofluorescence at scale. HeLa cells treated with NCS (1 *µ*g/ml, 5 min, followed by 15 min recovery) and stained for *γ*H2AX and phosphorylated Chk1, N=11,000 per condition. The scatter plot shows single-cell measurements, while the histogram and cumulative distribution show a shift in *γ*H2AX with no accompanying change in phosphorylated Chk1. Acquisition, segmentation and quantification proceeded without operator intervention.

### 5.2 Sustained observation of an unsynchronized population

The live-imaging application was used to test whether the system could sustain unattended observation long enough to capture transitions that occur at unpredictable times.

HeLa cells stably expressing PCNA-chromobody were treated with 4-nitroquinoline 1-oxide (4NQO), a UV-mimetic agent, and imaged every 20 minutes for 24 hours across many fields, with focus maintained automatically between time-points. PCNA localization reports both replication and repair, in that punctate signal in S phase reflects replication foci, while punctate signal outside S phase indicates sites of ongoing repair.

Cells were followed through division in both conditions. Control G1 daughters showed the diffuse PCNA signal expected outside S phase, while treated G1 daughters showed punctate PCNA (Fig. 6). The images are representative of the cells that were followed. The proportion of daughters showing puncta was not quantified and cell-cycle stage was assigned by manual inspection, so this is reported as an illustration of what sustained observation makes visible and not as a characterization of repair inheritance.

**Figure 6.**
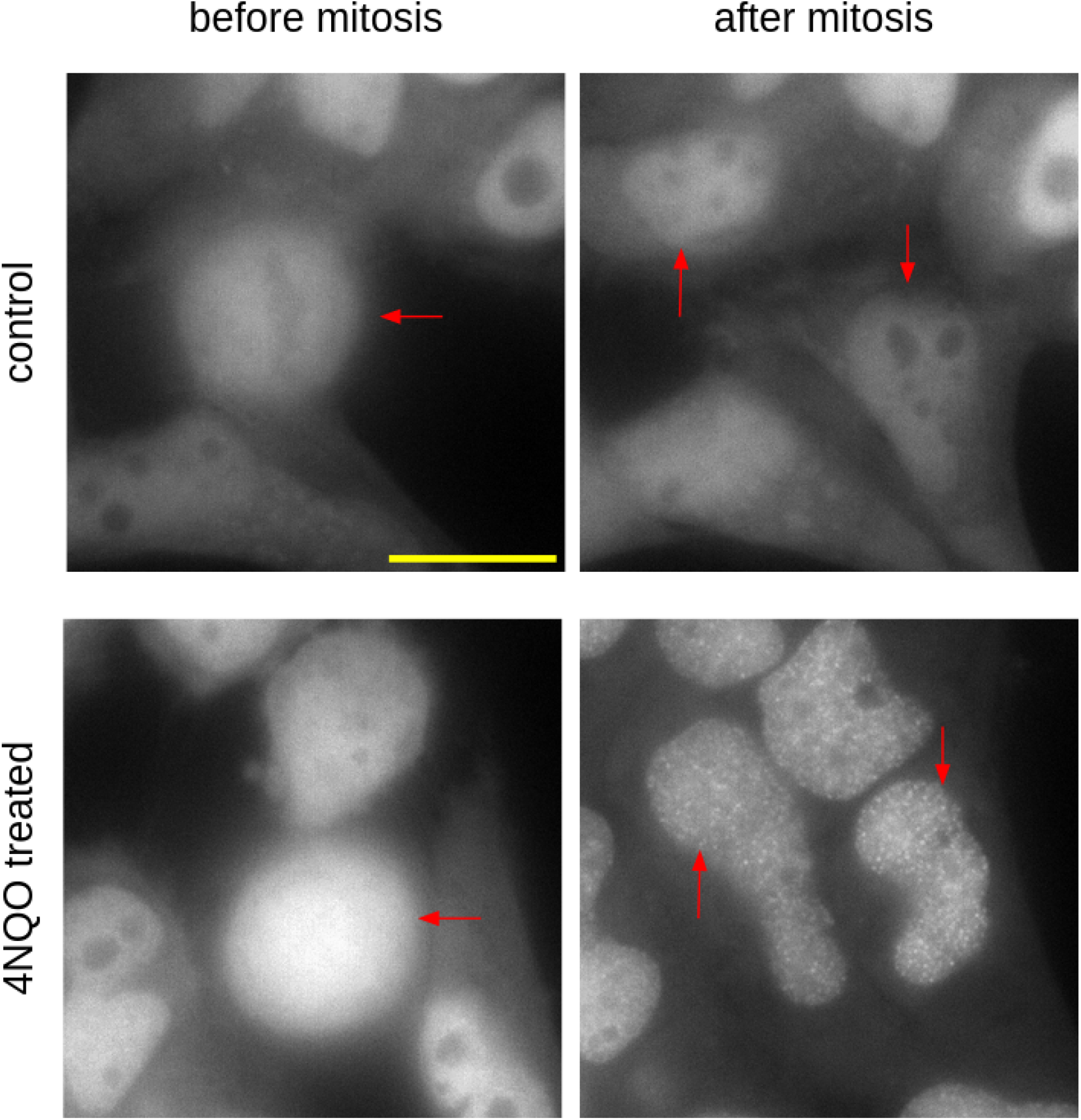
Sustained observation through cell division. HeLa cells expressing PCNA-chromobody were imaged every 20 minutes for 24 hours in control and 4NQO-treated conditions (1 *µ*g/ml, 15 min, followed by PBS wash), and cells were followed through division. Panels show one representative example from each condition, before and after mitosis. The control G1 daughter cells show the diffuse PCNA signal expected outside S phase, while the treated G1 daughters shown here have punctate PCNA. Arrows mark parent (left) and daughter cells (right). The proportion of daughters showing puncta was not quantified, and cell-cycle stage was assigned by manual inspection of morphology.

What the architecture provides is the observation itself. Following a cell through division in an unsynchronized population requires many fields to be revisited over many hours, with focus held between visits and no operator present. The alternative is to synchronize the culture chemically so that division can be anticipated, although the arresting agents used for this can themselves perturb the damage response being measured. Imaging enough cells for long enough removes the need to synchronize, and with it the confound that synchronization introduces.

## 6 Discussion

### 6.1 Generality of the classification

The classification in Table 1 can be applied independently of this system. Given a stack and a set of applications, each element can be placed by asking how it behaves across them. An element that is invariant with an implementation already available can be adopted, an element that is invariant with none available has to be built, an element that varies can be defined as an interface, and an element arising in a single application can be left as a local detail. The classification is testable, since adding an application in which a previously invariant element varies would show that the element had been misclassified.

This may address the adoption problem more directly than a further framework would. Capable systems already exist [Roos et al., 2024, Marin et al., 2024]. What has been difficult is that adding closed-loop capability has generally meant migrating to the acquisition stack that provides it. If most of the stack is already solved and only two representations are new, the same capability can be added to an installed system by changing what a run emits and what a plan consumes, without the acquisition system itself being replaced.

A second consequence of separable elements is that they can be worked on separately, and by different people (Fig. 7). A biologist who never writes a device adapter might define what counts as an object of interest, while an instrument developer might add hardware support without knowing what the resulting data will be used for. This is a design intention and not an observed outcome, since our own use of the system spanned all the elements, and how others divide the work has not been studied.

**Figure 7.**
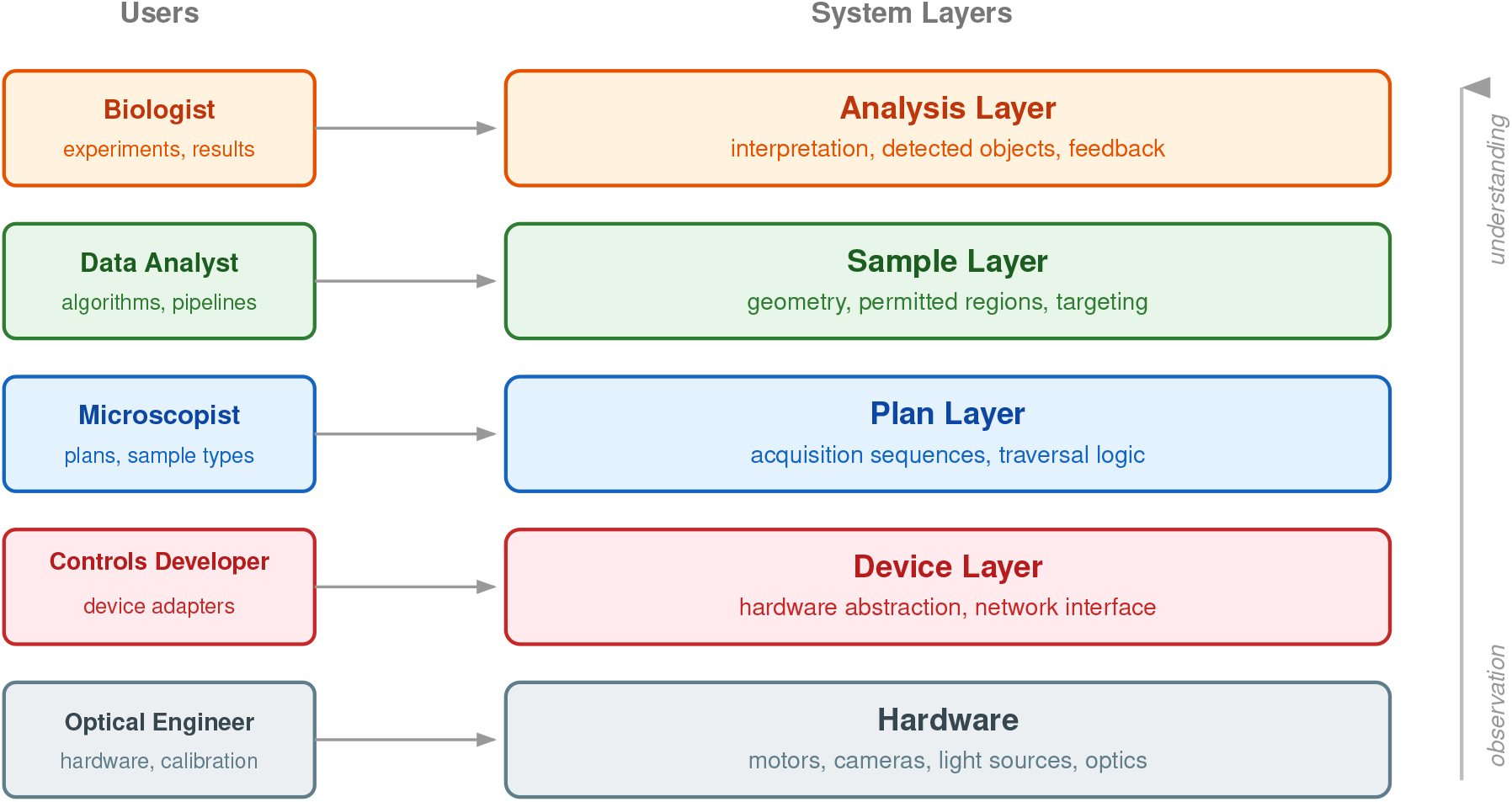
Elements as points of entry. Since each element is separable, work on it requires understanding only its interface to the elements adjacent to it. The vertical axis represents the transformation from raw observation to interpreted result as data moves upward. The division of labour shown here is the intent of the separation and not a measured pattern of use.

### 6.2 Targeting, an unabstracted element

The study identified one element that was not resolved here. Targeting, or the decision of where the next observation should be made, must be answered by every application, and applications answer it incompatibly. High-throughput immunofluorescence and time-lapse imaging both target a grid derived from the sample geometry, and this strategy is implemented using scanspec, which expresses the traversal as a composable specification that is then intersected with the permitted region of the sample. A fifth application, specified and partially implemented, targets clusters of previously detected objects instead, in which entities are grouped, each grouping is scored by how well it fits within a single field of view, and fields are then placed so as to maximize the number of objects imaged per acquisition. These two strategies share no parameters. The first derives positions from the specimen’s geometry before any data is acquired, while the second derives them from what has already been observed. By the criterion used here, targeting should therefore be treated as an interface, with the grid and the clustering strategies as two implementations of it. That is not what was built. The grid strategy is constructed inside the plan, and a traversal can only be supplied in advance as a geometric specification, which by construction cannot consume detections. Adopting scanspec supplied an implementation of one strategy and, in doing so, made it easy to leave the element itself unabstracted.

The fifth application was consequently not completed, and it is reported here for what it revealed about the stack. Targeting is also the element that must vary for a microscope to be adaptive and not merely automated, since it is the only element whose answer depends on data acquired during the run. A useful next contribution in this area can therefore be expected to be an abstraction for targeting.

### 6.3 Development history

The architecture reported here was not derived from a principle and then implemented. It is the outcome of building and rebuilding a working system over several years, during which most components were first written by hand and subsequently replaced once the concept they implemented came into focus. Sample representation appeared as an unimplemented specification roughly two years be-fore any code satisfied it. The development record, including the applications that were abandoned and the components that were discarded, is public in the repository history, with the earliest work preserved separately at https://github.com/ndsystems/deepthought-react.

### 6.4 Limitations

Four applications is a small breadth, and all four were run on wide-field instruments in one laboratory. Elements classified here as invariant might vary for modalities that were not tested, and light-sheet imaging, which couples illumination geometry to sample mounting, is the most likely to reclassify sample representation. The applications were selected because they were experimentally necessary to the biology being pursued, and their spread across demands in space, time, optics and control was recognized as useful during the work and not designed in advance. A study constructed deliberately to span modalities would test the classification more severely than this one does.

Two of the four applications, being anisotropy imaging and autonomous exposure, were built only far enough to exercise the elements of the stack, so the classification rests on two applications carried to completion and two carried partway. The biological findings reported here are consistent with existing understanding of the damage response and are presented as demonstrations of what the architecture makes routine, and not as a full characterization of either phenomenon. In the live-imaging experiment, the assignment of cells to cell-cycle stage involved manual inspection of morphology, and an automated criterion would be required to scale that analysis further.

### 6.5 Conclusion

Most elements of the microscopy software stack already have mature external implementations, and adopting them concentrates development effort on the elements that do not. Building a stack and driving several applications through it separates what has to be built from what can be adopted, and in this case it reduced the novel content to two representations, being the sample as a geometric object which plans traverse, and the run as a producer of detected objects. With both in place, the feedback path is ordinary control flow. A third requirement, targeting, is identified here and remains open.

## 7 Methods

### 7.1 Cell culture and maintenance

HeLa cells and HeLa cells stably expressing PCNA-chromobody were cultured in DMEM/F12 supplemented with 10% FBS (Gibco, 16000-044) and 1% PenStrep-glutamine (Gibco, 10378-016). Cells were maintained in T-25 flasks (Corning) at 37°C with 5% CO_2_ in a humidified incubator (Eppendorf Galaxy 170S). For imaging experiments, cells were plated on glass-bottom dishes (Genetix, 200350) 24 hours before experimentation.

### 7.2 DNA damage treatments

For DNA double-strand break induction, cells were treated with neocarzinostatin (NCS, Sigma, N9162) at 1 *µ*g/ml for 5 minutes followed by a 15-minute recovery period. For UV-mimetic damage, cells expressing PCNA-chromobody were treated with 4-Nitroquinoline 1-oxide (4NQO, Sigma) at 1 *µ*g/ml for 15 minutes, followed by PBS washes. Live-cell imaging was performed in FluoroBrite medium (Gibco, A18967-01) supplemented with 10% FBS and 1% PenStrep-glutamine.

### 7.3 Immunofluorescence

Cells were fixed with 4% paraformaldehyde (Sigma P6148) in PBS for 10 minutes at room temperature. After two PBS washes, cells were permeabilized using 0.3% Triton X-100 (Sigma, T8787) in PBS for 10 minutes. Following permeabilization and two PBS washes, samples were blocked with 5% BSA (Himedia TC545) in PBS for one hour. Primary antibodies were diluted in blocking solution and applied overnight at 4°C. After three PBS washes, samples were incubated with secondary antibodies in blocking solution for 2 hours at room temperature. Following three additional PBS washes, nuclei were stained with DAPI (1 *µ*g/ml in PBS) for 10 minutes. Samples were maintained in PBS for imaging unless otherwise specified.

### 7.4 Microscopy and image acquisition

Images were acquired using a 60x objective on an Olympus IX83 motorized inverted microscope (Tokyo, Japan) equipped with an Andor Zyla 4.2 scientific CMOS camera. Instrument access was provided by a server process exposing the micro-manager core interface over TCP/IP, configured using standard micro-manager device configuration files, with acquisition parameters defined through the framework described here.

Live-cell imaging was performed using an environmental chamber by Okolab UNO (Naples, Italy) maintaining 37°C, 5% CO_2_, and humidity. Time-lapse sequences were acquired at 20-minute intervals for 24 hours, with automated focus maintenance between timepoints. Multiple positions were imaged in parallel to maximize data collection while minimizing photobleaching and phototoxicity.

### 7.5 Image analysis and data processing

Image analysis was performed during acquisition. For fixed-cell analysis, nuclear regions were identified from DAPI images using Cellpose [Stringer et al., 2021] (version 0.6) with the nuclei model, a nominal object diameter of 100 pixels and a single input channel, run on GPU. Objects touching the field boundary were excluded. Intensity measurements for *γ*H2AX and phosphorylated Chk1 were extracted from each nuclear region using the DAPI-derived label as a common mask across channels, with local background subtraction applied to correct for imaging artifacts. Detected objects were associated with positions on the sample and accumulated during the run.

The system was run with micro-manager 2.0, ophyd 1.6.0, bluesky 1.6.7, databroker 1.2.0, napari 0.4.3, Cellpose 0.6 and scikit-image 0.18.1, with instrument access served over rpyc. For live-cell experiments, PCNA-chromobody signals were used to follow replication patterns and repair foci. Cells were tracked using nuclear signals, and division events were identified from characteristic changes in nuclear morphology, with the assignment of cells to cell-cycle stage confirmed by manual inspection. Analysis was performed using standard scientific Python libraries.

### 7.6 Statistical analysis

For comparisons between control and treated populations, Kolmogorov-Smirnov tests were used to evaluate differences in distributions, with the test statistic and associated p-value reported on the cumulative distribution plots.

### 7.7 Software availability

The system described here is a research testbed, developed as an instrument for studying the stack and not to be deployed as a product, and it can be read as a reference implementation of the two representations reported in this paper. The source, together with the development history including discarded components and abandoned applications, is available at https://github.com/ndsystems/deepthought, with the instrument access server at https://github.com/ndsystems/hard-link. The repository is archived at the state in which the results reported here were produced, and it depends on library versions contemporary with that work.

### 7.8 Data availability

The imaging datasets underlying the results reported here are held on local storage and are not currently deposited in a public repository. They are available from the corresponding author on reasonable request.

## 8 Competing interests

P.S.K. is affiliated with NDimensional Systems, which has an interest in microscope control software. The software reported here is released under an open licence and the authors declare no other competing interests.

## 9 Acknowledgements

We thank the members of the Mazumder lab at TIFR Hyderabad for their valuable discussions and support throughout this project. We are grateful to the Imaging and Flow Cytometry Facility (supported by DAE project # RTI 4007) at TIFR Hyderabad for their technical assistance. Special thanks to Jensun Ravichandran for providing important inputs during software development. We also thank the bluesky community for their work on scientific data acquisition software design, which shaped this project substantially.

## Notes

### Summary of Updates

Summary of changes (v2 to v3) FRAMING. Version 2 presented deepthought as an application of domain-driven design to microscopy software. This version reports the acquisition stack itself as an object of study. A minimal stack was built end to end, four applications were driven through it as test conditions (fixed-cell high-throughput immunofluorescence, live time-lapse imaging of an unsynchronized population, fluorescence anisotropy imaging, and autonomous focus and exposure), and each element of the stack was classified by how it varied across them. The layered structure is now presented as a result of that classification rather than as a starting premise. New sections give the method, the criterion used to individuate elements, and the full classification. FINDINGS. Two elements had no implementation available to adopt and were therefore built: a geometric representation of the sample that a plan can traverse, and a representation of a run that yields detected objects rather than images. Closed-loop operation follows from the second and required no additional machinery. A third element, targeting, or the choice of where to observe next, varies between applications, was never exposed as an interface, and is reported as an open requirement rather than as unfinished work. ADOPTED COMPONENTS. The externally sourced parts of the stack are now named with versions (micro-manager and MMCore, ophyd, bluesky, databroker, napari, Cellpose, scanspec, rpyc) and reported as an outcome of the classification rather than as implementation detail. Cellpose and napari are now cited. RESULTS. The sample-size comparison shown in version 2 has been removed. The underlying per-cell data is not currently accessible, and the comparison rested on single subsets at each sample size rather than repeated draws. A live-imaging result is added: cells expressing PCNA-chromobody were imaged every 20 minutes for 24 hours and followed through division in an unsynchronized population, without chemical synchronization. It is presented as a demonstration of sustained unattended observation. The proportion of daughter cells showing punctate PCNA was not quantified and cell-cycle stage was assigned by manual inspection, and both are stated in the text and the caption. AUTHORSHIP. S. Devadasan is added as an author for contributions to the software. OTHER. The title has been changed to reflect the reframing. Competing interests and data availability statements have been added. Figure annotations and placement have been corrected, and the software availability statement now describes the repository accurately as an archived research testbed.

